# Characterization of cellular wound resistance in the giant ciliate *Stentor coeruleus*

**DOI:** 10.1101/2025.06.23.661154

**Authors:** Rajorshi Paul, Ambika V. Nadkarni, Jana Sipkova, Justine E. Sato, Kevin S. Zhang, Vanessa Barone, Wallace F. Marshall, Sindy K.Y. Tang

## Abstract

Resistance to mechanical stress is essential for cells to prevent wounding and maintain structural integrity. This capability is especially critical for free-living single-celled organisms, which routinely encounter mechanical stress from their natural habitats. We investigated *Stentor coeruleus*, a single-celled ciliate known for its remarkable wound repair capacity, as a model for studying mechanical wound resistance. While previous work focused on wound repair in *Stentor*, the structures that enable it to resist wounding remain poorly understood. We characterized how *Stentor* resisted mechanical stress during transit through a microfluidic constriction. Using high-speed imaging, we tracked the transit dynamics of the cells and linked them to wounding outcomes. Larger cells experienced longer transit times in the constriction and were more prone to rupture, often failing to recover shape due to membrane rupture and loss of cytoplasm. To elucidate the role of the *Stentor* cytoskeleton, we performed drug-mediated disruption of KM fibers, which are microtubule bundles in the *Stentor* cytoskeleton. Drug-treated cells exhibited an increased likelihood of membrane rupture at the constriction, implicating KM fibers in wound resistance. To investigate the resistance of *Stentor* cells to hydrodynamic stress, we injected the cells at increasing flow rates through the constriction. Interestingly, cells were more resistant to larger hydrodynamic stresses up to a threshold, potentially due to shear-thinning of the cytoplasm. Together, these results suggest that *Stentor* relies on both cytoskeletal architecture and cytoplasmic rheology to withstand mechanical stress, offering insights into cellular strategies for wound resistance in the absence of rigid extracellular structures.

## Introduction

Mechanical forces are integral to cellular function. For example, mechanotransduction, which involves sensing mechanical forces in a cell’s microenvironment, is critical for promoting^1^ or inhibiting^2^ cell proliferation, volume regulation,^3^ activation,^4^ and differentiation.^5–7^ Some cells routinely experience mechanical stress as part of their normal function—for example, endothelial cells in blood vessels during blood flow,^8^ chondrocytes under continuous mechanical loading,^9^ and muscle cells including cardiac myocytes during contraction and stretching.^10,11^ Because cell membranes are thin and fragile, the ability to resist mechanical wounds is essential for the cells to maintain structural integrity and prevent recurrent wounds. Wound resistance is especially critical for single-celled free-living organisms. In their natural habitats, these organisms can be exposed to substantial mechanical stresses from predation or other environmental factors. To survive, they must evolve mechanisms to resist wounds. Many cells have adaptations such as cell walls, exoskeletons, cysts, or spores.^12–14^ Perhaps more interesting are cells that do not have rigid outer shells, such as ciliate protozoa, which must rely on internal cellular structures to provide wound resistance.

We focused on *Stentor coeruleus* as our model system to understand how cells without external defense structures can resist mechanical wounding. *Stentor* is a single-celled ciliate found in freshwater ecosystems and is known for its ability to repair drastic wounds.^15–17^ Although we and others have investigated wound healing and regeneration in *Stentor*,^16–33^ the extent to which *Stentor* can resist mechanical wounds, and the role of cellular structures that enable this wound resistance, remain largely unknown. Previous studies in other cell types have shown that components of the cytoskeleton, such as microtubules and actin filaments, can bear mechanical load and resist compression,^34^ as observed in contracting cardiomyocytes^35^ and myoblasts.^36,37^ The collective behavior of the cytoskeletal network also gives rise to viscoelastic properties, allowing cells to regain their original shape following mechanical deformation.^38^

Based on these findings, we propose that the cytoskeleton in *Stentor* also contributes to wound resistance. Electron microscopy of *Stentor* has revealed fine morphological details of two main elements of the *Stentor* cytoskeleton, the KM fibers and the myoneme cytoskeleton.^39^ KM fibers are bundles of prominent, highly stable, acetylated microtubules that run down the length of the cortex in *Stentor*. KM fibers are closely apposed to the outer pellicle, a membrane of flattened alveolar sacs and the oral and body cilia. Each cell has many parallel KM fibers, spaced approximately 3-5 μm apart.^40^ Myonemes, present underneath the KM fibers, are made up of 2-4 nm filaments of a caltractin-like protein.^40,41^ Myonemes work antagonistically with the KM fibers, providing contractile forces.^40^ KM fibers and myonemes are connected by fibers of unknown composition.^40^ Other proteins and organelles could participate in the formation of cytoskeletal networks, however, their molecular identity and configuration remain undefined.

In this work, we subject *Stentor* cells to mechanical stress in a microfluidic constriction and characterize the conditions under which wounding occurs. Although microfluidic constrictions have been used for mechanical characterization of cells,^42–47^ this geometry has not been used for studying wound resistance in cells. By pharmacologically weakening the microtubule network and systematically varying the applied hydrodynamic stress, we identify factors that contribute to wound resistance in *Stentor*.

## Materials and methods

### *Stentor* cell culture

*Stentor coeruleus* were cultured in Pasteurized Spring Water (PSW) (132458, Carolina Biological Supplies) as previously described.^16–18^ We maintained two cell cultures and the feeding schedule was staggered so that each culture was fed *Chlamydomonas rheinhardtii* (∼0.2 μL per *Stentor* cell) once every two days. Prior to each experiment, healthy adult cells (∼200 - 400 μm in diameter and dark green in color) were retrieved from one of the cultures 36 – 48 hours post feeding. The cell retrieval was done by pipetting under a stereoscope into a 2 mL centrifuge tube. The cells were then washed once with fresh PSW.

### Microfluidic constriction design, fabrication and wounding experiments

The master mold of the microfluidic device was fabricated in SU-8 on a silicon wafer using standard photolithography as described previously.^16–18^ The height of the master was measured with a profilometer to be 120 μm. The channel height was chosen such that the *Stentor* cells, which were 200–400 μm in size, were compressed and fully occupied the height of the microfluidic device. Poly(dimethylsiloxane) (PDMS) (SYLGARD-184^TM^, Dow Corning) was then cured from the original master following standard soft lithography procedures and bonded to a glass slide to form the final device. The device was left overnight at 65 °C in an oven to strengthen the adhesion between PDMS and glass and to make the channel hydrophobic. Prior to use, at least 1 mL of cell media was injected to clean the channels. Channels were washed from the outlet with cell media between separate experiments and discarded if cell debris could not be removed from the channel. All microfluidic channels and tubing for cell injection were passivated using 3% (w/v) F-68 polymer solution for 1 hour to reduce friction on all surfaces that the cells encountered. This surface treatment was done to prevent injury of cells prior to the cells reaching the constriction.

The microfluidic channels used in this work are shown in ESI Fig. S1. The straight part of the channel leading to and away from the constriction was 250 μm in width and was designed to have a 15° ramp to the constriction (as indicated in Fig. S1A, ESI). The constriction tapered to a point and then diverged. We varied the constriction width, defined at the narrowest point of the constriction, across 30, 35, 40, 45, 50, 70, 85 and 100 μm. The length of the constriction starting from the converging section to the end of the diverging section varied from 807 μm (for the 100 μm constriction) to 1036 μm (for the 30 μm constriction). All cell velocities reported in this work were measured in the straight part of the channel upstream of the constriction. We observed that when the flow rate was higher than 10 mL/h (Fig. S1A, ESI), some cells appeared to get wounded because of the long transit through the channel at high shear rates. To circumvent this issue, we designed an additional subsidiary channel to apply a sheath flow (Fig. S1B, ESI). In the high velocity channels, the cells were injected at the primary inlet at 10 mL/h, and a sheath flow of 13 mL/h was added through the secondary inlet to generate a final flow rate of 23 mL/h at 600 μm (∼1–2 cell body lengths) before the constriction entry, thereby preventing the cells from injury prior to entering the constriction.

To flow cells into a microfluidic channel, a cell suspension was first obtained by collecting cells from the centrifuge tube into tubing attached to a syringe filled with PSW. The inner diameter of the tubing (762 μm) was large enough so that cells did not get wounded while flowing through the tubing. The syringe was driven by a syringe pump (Chemyx Fusion 101/Chemyx Fusion 4000/ Kent Scientific GenieTouch) and the end of the tubing was attached to the inlet of the microfluidic device. Finally, the cells were injected into the microfluidic channel by applying a desired flow rate through the syringe pump. The flow rates were 2, 4, 7, 10, and 23 mL/h, corresponding to an average velocity of 0.02, 0.04, 0.06, 0.09, and 0.21 m/s, respectively in the straight part of the channel outside the constriction.

The experiments were imaged with the Vision Research Phantom v7.3 or v341 high speed cameras mounted on an inverted microscope (Nikon Eclipse Ti-U) stage with a 4x Nikon objective lens (numerical aperture 0.1, working distance 30 mm) The frame rate for imaging varied from 450 frames per second (for flow rates of 2 and 4 mL/h) to 5000 frames per second (for flow rates higher than 4 mL/h).

### Immunostaining of acetylated tubulin to visualize KM fibers

To visualize KM fibers in cells, we performed immunofluorescence for acetylated tubulin. To visualize the effect of cytoskeletal drugs on the KM fiber organization, we first treated the cells with drugs (or DMSO as the control - method described in the next section), and then stained the cells. To visualize the cytoskeletal disruption due to the microfluidic constriction, we first flowed cells (treated or untreated) through the microfluidic constriction at 4 mL/h. The cells were collected at the end of a 4 cm long tubing placed at the outlet of the microfluidic device at ∼16 seconds after the cell exited the constriction. The control condition for this case involved flowing untreated cells through a tubing at the same flow rate and collecting the cells at the end of the tubing.

To begin the staining process, we fixed 250 μL of cells in 1000 μL of ice-cold methanol in a 2 mL plastic tube (16466-060, VWR) as previously described.^17,18^ Briefly, cells were incubated in methanol at -20 °C overnight. Cells were then rehydrated with 1:1 PBS:methanol, and 1x PBS pH 7.4. Cells were blocked with 2% BSA + 0.1% Triton X-100 in PBS and stained with 1:250 mouse anti-acetylated tubulin primary antibody (T7451-200UL, Sigma Life Sciences) followed by 3 washes in 1x PBS. The washes were done by gently pipetting as much of the supernatant liquid as possible without disturbing the fixed cells and adding fresh PBS. We waited for the fixed cells to settle down before the next wash. Cells were stained with anti-mouse secondary antibody (1:500). (SAB4600388-125UL, Fluka Analytics).

Cells were imaged using the Zeiss LSM 780 laser scanning microscope, using a 20X (NA = 0.8) objective at an excitation wavelength of 488 nm, and a broad emission filter matching the spectra of Alexa Fluor 488.

### Drug treatment experiments

500 μL of culture media (i.e., PSW) containing ∼30 cells were collected in a 2 mL round bottomed centrifuge tube. The cells were then washed by adding 500 μL of fresh PSW to the tube and carefully removing 500 μL of liquid carefully without removing the cells. PSW was added gently to ensure that we did not wound the cells. In a separate 2 mL round bottomed centrifuge tube, we prepared a 50 μM solution of the drug which was either nocodazole (Sigma Aldrich, Product number M1404) or taxol (Tocris, Catalog number 1097) inside a chemical safety hood. To do this, we first brought a 50 mM aliquot of the drug solution in dimethyl sulfoxide (DMSO) from -70 °C to room temperature. We then added 0.5 μL of the 50 mM drug solution to 500 μL of PSW in a 2 mL centrifuge tube to prepare a final drug concentration of 50 μM. Next, we added 500 μL of the 50 μM drug solution to 500 μL of the cells previously collected so that the final concentration of the drug for treatment was 25 μM. The control condition (0.05% DMSO in PSW) was prepared in the same way, by replacing 50 mM of drug solution with DMSO. The cells were treated for 90 mins at room temperature in the dark. After incubation, the solutions were washed with PSW before taking the samples out of the safety hood in preparation for the experiments.

### Image processing and data analysis pipeline

The video data collected from experiments were processed using the OpenCV package in Python. Custom image processing codes were used to identify the cell and track its trajectory along the channel. The kinematic data was used to extract parameters of interest. We used a pixel intensity-based threshold for the purpose of identifying the cell boundary from its surroundings. Under the lighting conditions in our experiments, the cells had a lower intensity than the surrounding microfluidic channel. To minimize the effects of non-uniform illumination, we subtracted a reference frame not containing a cell which helped highlight the cell from the surroundings even more. In the case of membrane rupture, the spilled content appeared to have similar pixel intensity as that of the intact cell, and therefore, was incorporated within the cell boundary.

The effective cell size, D, was estimated upstream of the constriction based on the projected cell area (i.e., the area enclosed by the cell boundary), A, i.e., D=√(4A/π).

The boundary of the cell was used to fit a bounding box around the cell. The sides of the bounding box aligned with the orientation of the image frame and therefore the microfluidic channel. The aspect ratio of the cell was measured as the ratio of the sides of the bounding box as indicated in Fig. 1B.

**Fig. 1.**
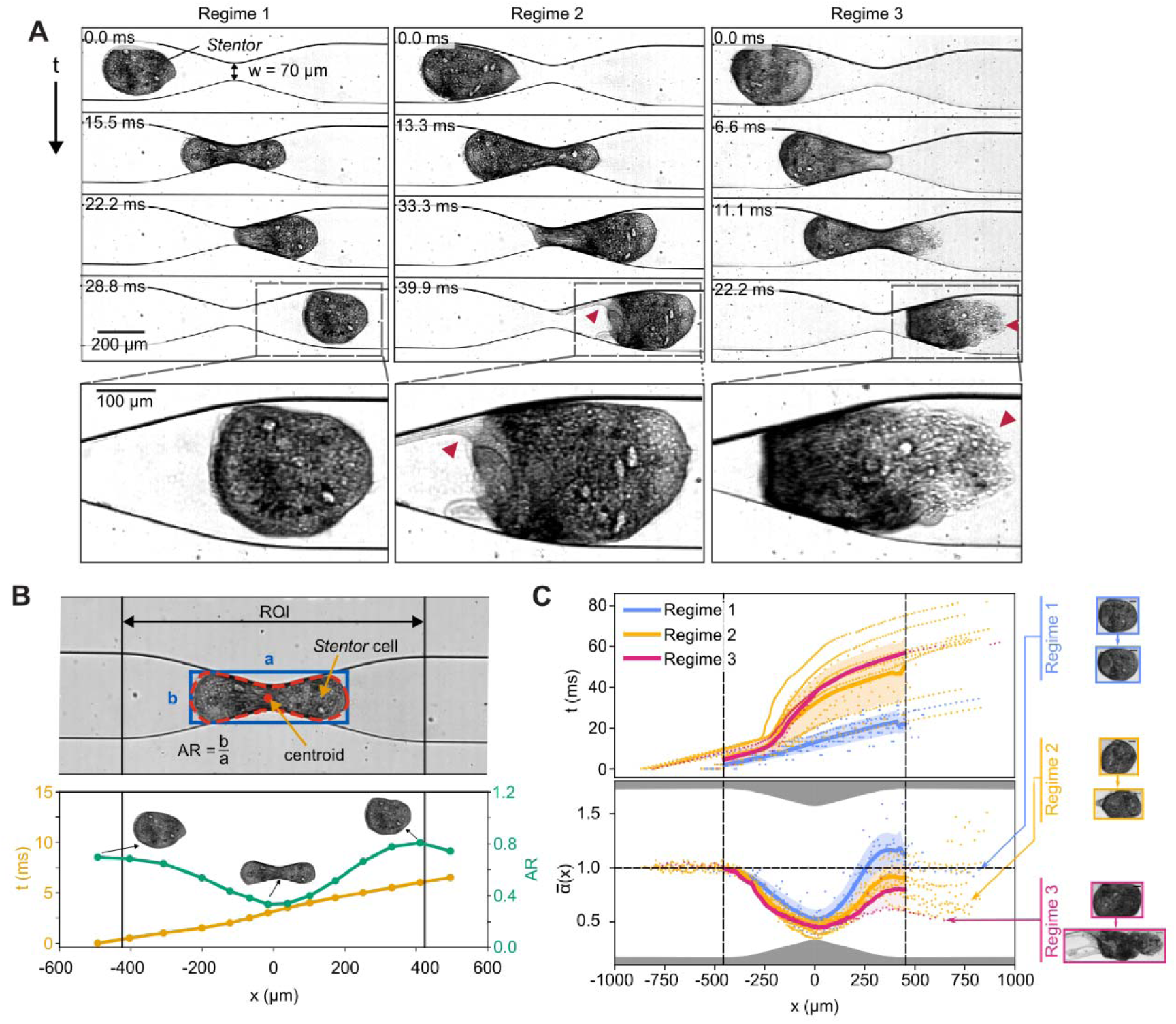
Characterization of wounding outcomes. **A.** *Stentor* cells injected into a microfluidic constriction showed three regimes of wounding outcomes. The cells were injected at 4 mL/h through a 70 μm constriction. The red arrows indicate sites of disruption in the plasma membrane. **B. Top:** Parameters extracted using image processing. **Bottom:** The trajectory of a cell, flowing through a 70 μm constriction at 4 mL/h, as a function of the position x along the microfluidic channel. The center of the constriction is at x = 0. The yellow curve shows the time and the green curve shows the variation of the cell aspect ratio (AR). The images of the cell at three different positions are shown. **C.** A compilation of all *Stentor* trajectories that flowed through a 70 μm constriction at 4 mL/h with time series data in the top panel and aspect ratio variation 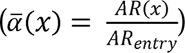 in the bottom panel. The trajectories are demarcated by the regime of wounding outcomes. The line and the shading indicate the mean and standard deviation for the trajectories for a given regime of wounding outcome within the region of interest. The insets show the cell deformation at the entry and exit for regimes 1 (blue), 2 (yellow) and 3 (red). The corresponding bounding boxes for the cells are shown. The scale bars represent 50 μm.

The transit time t is measured from the frame when the cell entered the region of interest (ROI, see Fig. 1B) until the cell completely exited the ROI. t_0_ is the ideal time taken by a fluid of the same volume as the cell to transit through the ROI and is estimated as 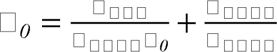 (see ESI Fig. S2 for details). Here, w_0_ = 250 μm is the width of the straight part of the channel, A_ROI_ is the area enclosed by the ROI, l_cell_ is the length of the cell estimated as the length of the bounding box, and v_cell_ is the cell velocity. l_cell_ and v_cell_ are measured prior to the cell entering the ROI.

For labeling the experimental data, we examined the videos and classified the data into three regimes (details in the Results and Discussions section). We discarded data points when: debris was attached to the cells, cell debris clogged the constriction, multiple cells attached together or very close together, the cells were very small, or the cells were already wounded prior to the constriction, which can sometimes occur during pipetting. These discarded data accounted for ∼10% of all the data that were collected (varying from 4% to 15% within the different flow velocity data sets). Additionally, we discarded data where there was uncertainty with the correct label assignment, accounting for ∼3% of all the data that were collected. These uncertainties were caused by factors such as inadequate resolution of the cell membrane, insufficient field of view downstream of the constriction, the orientation of the cell (making it challenging to classify an outcome) and inadequate illumination. Discarding these data did not impact the overall interpretation of our results.

For plotting cumulative probability distribution (Fig. 4B), the ranges of the cortex capillary numbers Ca_c_ were allocated by distributing the overall range of the capillary numbers into 5 bins based on equal logarithmic spacing. For estimating the cumulative distribution for each capillary number range, the data was distributed into bins containing approximately equal number of data points based on the D* values. The value of D* for each bin was estimated by averaging the maximum and minimum D* values for the bin.

### Rheometry

Rheometry was performed to characterize the shear rate dependent rheological behavior of the *Stentor* cells. We used a TA ARES G2 rheometer with a 25 mm parallel plate for the characterization. For the characterization, we loaded 50 μL of our sample onto the platform and brought the parallel plate to the desired gap. In our measurements, we used a 100 μm gap between the plates so that the vertical confinement of the cells is similar to that inside the microfluidic channel and to minimize the sample volume requirement. The 100 μm gap also ensured that there was one layer of cells (cell size ∼ 200 – 400 μm) between the plates. We trimmed any excess sample from around the plates while ensuring that the gap was completely filled with the sample. During the measurement, we increased the shear rate from 1 s^-1^ to 5000 s^-1^ while recording 10 data points per decade. All measurements were performed at 22°C.

We characterized three samples: PSW without cells, PSW with ∼6% volume fraction of cells, and PSW with ∼12% volume fraction of cells. Cell volume fraction, here is defined as the ratio of the cell volume to the volume of PSW. The cell samples were prepared by collecting cells from the main culture into 2 mL microcentrifuge tubes and successively removing excess liquid to increase the volume fraction. 6% and 12% volume fractions of cells corresponded to 300 and 600 cells respectively in 50 μL of medium (PSW) with each cell being estimated to be ∼1 nL in volume.

### Micropipette aspiration

#### Cell preparation

*Stentor* cells were left to adhere to 60 mm glass-bottomed dishes (P60G-1.5- 20-F, MatTek) in Pasteurized Spring Water (PSW) for ∼18 hours at 22°C. To reduce cell movement during the experiment, a 5 min salt shock with 7.5% filtered sea water (salinity: 36 ppt) in PSW was applied. Cells were then maintained in PSW for the remainder of the experiment. For the nocodazole condition, the drug treatment was performed prior to the salt shock. Only healthy, viable *Stentor* cells, including those that visibly responded to mechanical stimulation, were selected for measurements.

#### Micropipette aspiration setup

Micropipette aspiration, as described previously,^48,49^ was used to measure cellular surface (cortical) tension. In *Stentor*, the cytoskeletal fibers, both (KM fibers and myonemes) are associated closely with the cell membrane.^40^ Therefore, the micropipette aspiration is expected to measure the cortical tension in *Stentor*. A 20 μm diameter blunt glass micropipette (VICbl-20-30-600-55, BioMedical Instruments) was passivated in heat-inactivated FBS (F0926-100ML, Sigma) for 10 min at room temperature (18°C) and then filled with PSW. The micropipette was mounted on a micromanipulator (Transferman 4r, Eppendorf) and connected to a Flow EZ™ pressure controller and LineUp™ LINK module (Fluigent), providing pressure control with 0.7 Pa resolution. Micropipette movement and pressure were controlled using Fluigent OxyGEN software. The setup was mounted on an inverted Zeiss LSM980 microscope with a ×20/0.8 air Plan Apochromat objective, and imaging was handled via Zeiss ZEN software. All measurements were conducted at room temperature (18°C).

#### Measurements

Calibration of the output pressure was performed using a ∼5 μm bead (CFP- 5045-2, Spherotech). The correct pressure was found when the bead remained stationary in the micropipette for ∼30 sec i.e. no flow was observed. Calibration was performed at the start of measurements in each dish. To measure cortical tension, the micropipette was positioned at a free surface of a stationary *Stentor* cell (typically near the oral apparatus). The pressure was then increased in 10 Pa steps until the cell protrusion aspirated into the pipette was the radius of the micropipette (R_p_) and remained stable for at least 20 sec (steady state). Using the Young-Laplace law, cortical tension *γ* was calculated as: 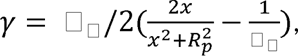 where P is the pressure used to deform the cell of radius R_c_, and *x* is the cell protrusion length in the micropipette.^50^ The first term in the expression refers to the radius of the curvature of the cell protrusion, when x ≤ R_p_. The negative pressure was then gradually reduced to release the cell. Each measurement lasted ∼1-3 min, and all measurements per dish were completed within 40 min to 1.5 hours after the salt shock.

#### Data analysis

Fiji ^51^ was used to measure the radius of the micropipette (R_p_) and the size of the deformation of the *Stentor* cell (*x*) using the line tool. A circle was fit at the site of the measurement to measure the radius of the local curvature of the *Stentor* (R_c_). The cortical tension was then calculated in Excel as described above. We discarded measurements where the length of the cell protrusion exceeded the micropipette radius by over 10%. Statistical analysis and plotting were performed in R using a Mann–Whitney U-test.

### Statistical analysis

Significance tests were performed in OriginPro 2024b using the two-sample t-test. For all comparisons, we performed Welch correction to obtain a more accurate estimate of the p values by assuming unequal variance between the datasets. For establishing statistical significance, we set the threshold for p value to be smaller than 0.05. We modelled the probability of rupture wounds (i.e., regime 3 outcome, defined in the next section) as a function of D*, the dimensionless cell diameter, by using the sigmoid function 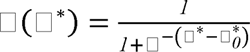 with the fitting parameter 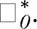 This approach was based on logistic regression used commonly in binary classification problems.^52^

To draw the decision boundary denoting the rupture (regime 3) probability as a function of the features and D*, we used the scikit-learn package in Python and performed a logistic regression with second-degree polynomial feature expansion on standardized input features D* and log_10_(Ca_c_) (Ca_c_: the cortex capillary number, defined in the text) on the binary outcomes (1 for regime 3 outcomes, 0 otherwise). The decision boundary was plotted corresponding to a predicted class probability of 0.10.

## Results and Discussions

### A microfluidic constriction to study mechanical stress-mediated wounding

To investigate the effect of mechanical stress on *Stentor* cells, we injected a dilute suspension of cells into a microfluidic constriction at a constant flow rate. The flow rate was chosen to ensure that the cells did not get wounded in the straight part of the channel. On entering the converging section of the channel into the constriction, the cell was exposed to mechanical stresses which caused the cell to deform. This deformation was driven by a combination of compressive stress from the converging channel, tensile stress due to the velocity difference across the constriction, and shear stress from the channel walls and fluid viscosity. The deformation caused by these mechanical stresses can lead to the disruption of the cell membrane.

We classified the outcome of cell deformation using high-speed brightfield imaging. We chose this method because it allowed us to capture the fast dynamics of the cells (ranging from 5 to 300 ms) and match kinematic information, such as transit time and deformation characteristics, to the corresponding cell deformation outcome. Other methods such as confocal imaging and assays using membrane-impermeable dyes are too slow to capture the deformation and membrane disruption dynamics. We classified the outcome of cell deformation into three regimes based on the degree of disruption to the cell membrane. Regime 1 refers to the outcome where no disruption was observable to the cell membrane within the field of view (Fig 1A, left panels). The membrane appeared to be intact before and after the cell transited from the constriction. Regime 3 refers to rupture wounds that led to the spilling out of the cytoplasmic contents from the cell (Fig. 1A, right panels). Cell rupture typically occurred on the downstream side of the cell on exit from the constriction. This observation is consistent with prior models on capsule transit through a constriction, where membrane deformation and stretching generates high stresses near the capsule tip.^53^ Regime 2 refers to any outcome where we observed some disruption in the cell membrane (e.g., shearing of the membrane and membrane blebs) without membrane rupture and spillage of cell cytoplasm (Fig. 1A, center panels; Fig. S3, ESI).

Fig. 1B outlines the image analysis pipeline for obtaining kinematic data of the cell as it transits through the constriction. The region of interest (ROI) encapsulates the converging and the diverging sections of the microfluidic constriction. The ROI is 900 μm in length and is centered around the center of the constriction (x = 0; x is the horizontal axis). We identified the boundary of the cell (shown in red in Fig. 1B (top panel)) and estimated the centroid and the deformation of the cell. Cell deformation was quantified using the aspect ratio (also referred to as “AR”) of the cell which was defined as the ratio of the sides of the bounding box (shown in blue in Fig. 1B (top panel)). The bottom panel of Fig. 1B shows the transit time t and the aspect ratio AR of a representative cell (with regime 1 outcome) as it traversed through the constriction in the x direction.

Fig. 1C shows variations in the kinematics of the cells for the three outcome regimes, for cells injected through a constriction with a width of 70 μm at a flow rate of 4 mL/h. At this constriction width, all three regimes were observed. We plotted the transit time t and the normalized aspect ratio 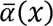 of the cells as a function of their position x through the constriction. Here, 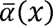 is defined as 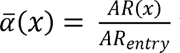 where AR(x) is the aspect ratio of the cell at position x, and AR_entry_ is the aspect ratio of the cell prior to its entry to the constriction (cell centroid was at x < -450 μm). Fig. 1C shows that cells in regime 1 (i.e., unwounded cells) transited through the constriction in shorter times than those in regimes 2 and 3 (i.e., outcomes with membrane disruptions). Upon exiting the constriction, cells in regime 1 became more prolate 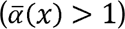 than cells in regimes 2 and 3. The prolate shape of the cells in regime 1 is attributed to the difference in velocity between the upstream and downstream sides of the cells as they exited the constriction. Similar to prior observations in droplets and capsules exiting a constriction, this velocity profile caused the cells to compress along the x direction and elongate in the perpendicular direction leading to an increase in the aspect ratio.^54–56^ In comparison, cells in regimes 2 and 3 had 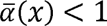 even after exiting the constriction, indicating that these cells retained their deformation in part due to the irreversible damage to the membrane/cortex and/or the spilling of the cytoplasm.

### Mapping the parameter space of cellular wound resistance

Given the geometry of the constriction, the width of the constriction and the size of the cell are expected to determine cell deformation and their wounding regime.^57,58^ At a constant flow rate (4 mL/h), narrower constrictions led to more severe cell wounding as indicated by more extensive breaks in the KM fiber network (Fig. 2A, red arrows). Because the cells used in our experiments had effective cell diameter D ranging between ∼ 190 μm and ∼ 410 μm, we defined a dimensionless cell diameter D* as the effective cell diameter D normalized against the constriction width w to compare results across cell sizes and constriction widths. Further, we defined a dimensionless transit time t* as a measure of the actual time t taken by the cell to transit through the ROI normalized against a characteristic timescale t_0_ for the fluid in the continuous phase with the same volume as the cell to transit through the ROI (see details in Fig. S2, ESI).

**Fig. 2.**
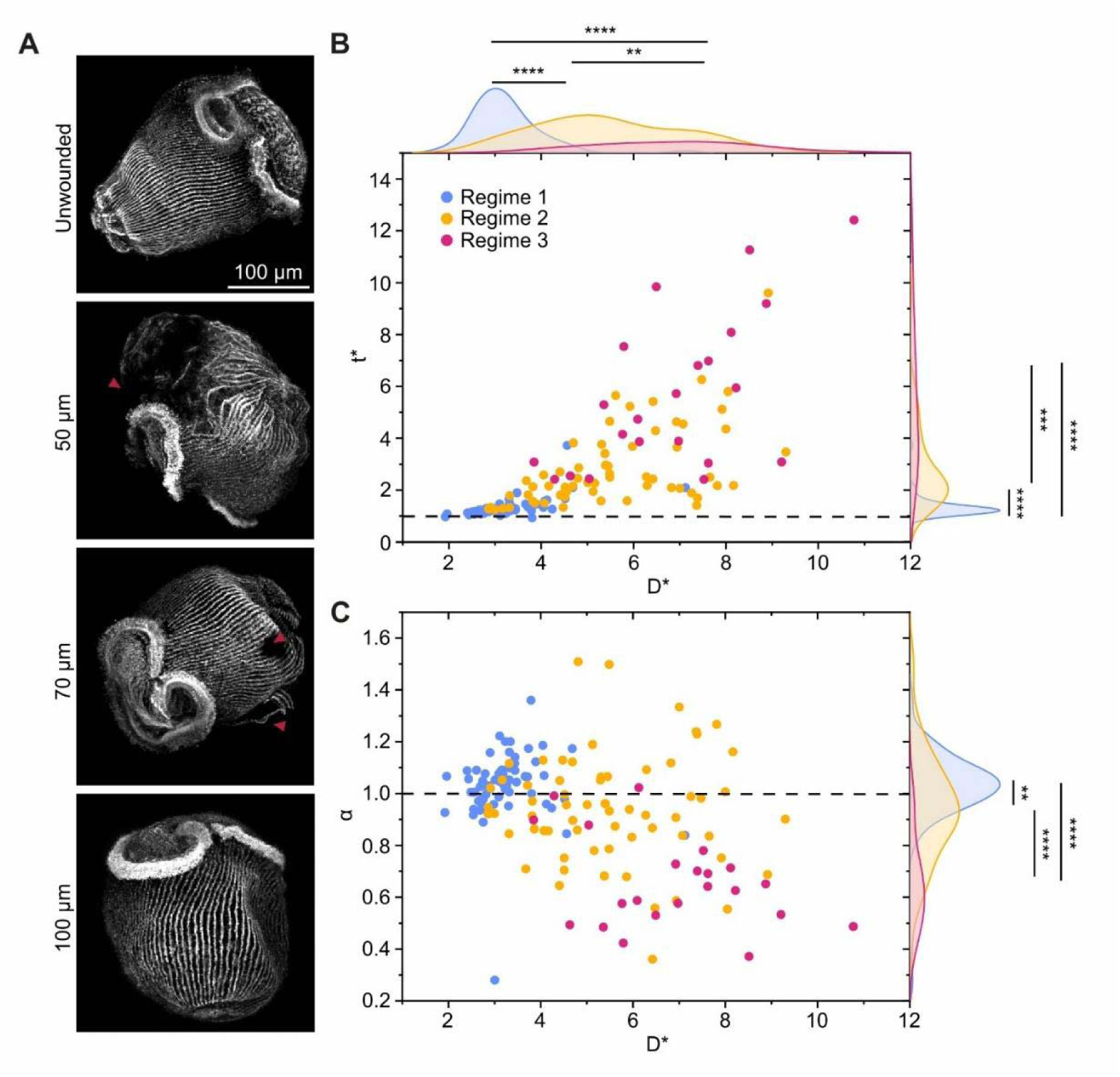
Relationship between cell size and wounding characteristics of *Stentor*. **A.** Immunofluorescence images of *Stentor* cells stained against acetylated tubulin show disruption of the KM fiber network after being injected through varying constriction widths. The red arrows indicate sites of KM fiber disruptions. The top panel depicts an unwounded cell that did not flow through the microfluidic constriction. **B.** Plot showing the dimensionless transit time t* and the dimensionless cell size D* for the three wounding regimes. The distributions of the data points for the three regimes for t* and D* show statistically significant differences. **C.** Plot showing the dimensionless parameter 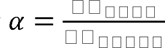 and the dimensionless cell size D* for the three wounding regimes. The distributions of the data points for α show statistically significant differences for the three wounding regimes.

Fig. 2B shows that as D* increased, the cell deformation outcomes transitioned from regime 1 to regime 3, with a corresponding increase in t*. In other words, larger cells (D* > ∼4) took longer to transit through the constriction and were more likely to be ruptured. The longer transit times occurred due to the occlusion of the constriction channel by large cells. Because we used a syringe pump to drive the flow at a constant flow rate, the occlusion subsequently raised the pressure differential across the constriction. This effect could also be seen in the flow of a capsule through a constriction.^54,59^ The increased mechanical stress increased the likelihood of membrane ruptures (i.e., regime 3 outcomes). An occluded cell was able to eventually transit through the constriction when the pressure gradient exceeded the forces exerted on the cell from the channel wall.

Next, we investigated the ability of the cells to recover their shape after transiting through the constriction. This shape recovery was measured by the parameter 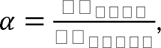 which compares the aspect ratio of the cells at the exit of the constriction to that on entry. Fig. 2C shows that as D* increased and as the cells transitioned from regime 1 to regime 3, *α* decreased from a median value of 1.04 (regime 1) to 0.93 (regime 2), and to 0.63 (regime 3). Consistent with Fig. 1C, these results indicate that the disruption of the membrane, along with the loss of cytoplasm, impeded the cell’s ability to maintain its structural integrity and recover its original shape.

### Cytoskeletal microtubule depolymerization reduces wound resistance

To assess the contribution of cortical KM fibers to cellular wound resistance, we treated *Stentor* cells with nocodazole. In mammalian cells, nocodazole is known to bind with tubulin dimers creating conformational changes, thereby inhibiting microtubule assembly^60^ and leading to the subsequent depolymerization of microtubules.^61^ Treatment of *Stentor* cells with nocodazole appeared to create alterations in the morphology of the KM fibers such as bunching, buckling and irregular puncta (Fig. S4). However, the fibers remained continuous.

To quantify the effect of nocodazole on the mechanical property of the *Stentor* cortex, we performed micropipette aspiration on the cells (Fig. 3A). The measured median cortical tension in control cells was ∼ 182 pN/μm, which is on the same order of magnitude as several other cell types including HeLa cells,^62,63^ mouse fibroblasts^64^ and mouse embryo.^48^ Notably, the cortical tension in cells treated with nocodazole (∼ 112 pN/μm) was nearly half of that in control cells (p = 0.023) (Fig. 3B). This result is consistent with the expected mode of action of nocodazole, which disrupts microtubule polymerization and weakens the KM fiber network. The cortical tension is often positively correlated with the elastic modulus of the cell cortex^65,66^ and serves as an indicator of the cortical stiffness. The reduced cortical tension may therefore reflect a reduced cortical stiffness in nocodazole-treated cells.

**Fig. 3.**
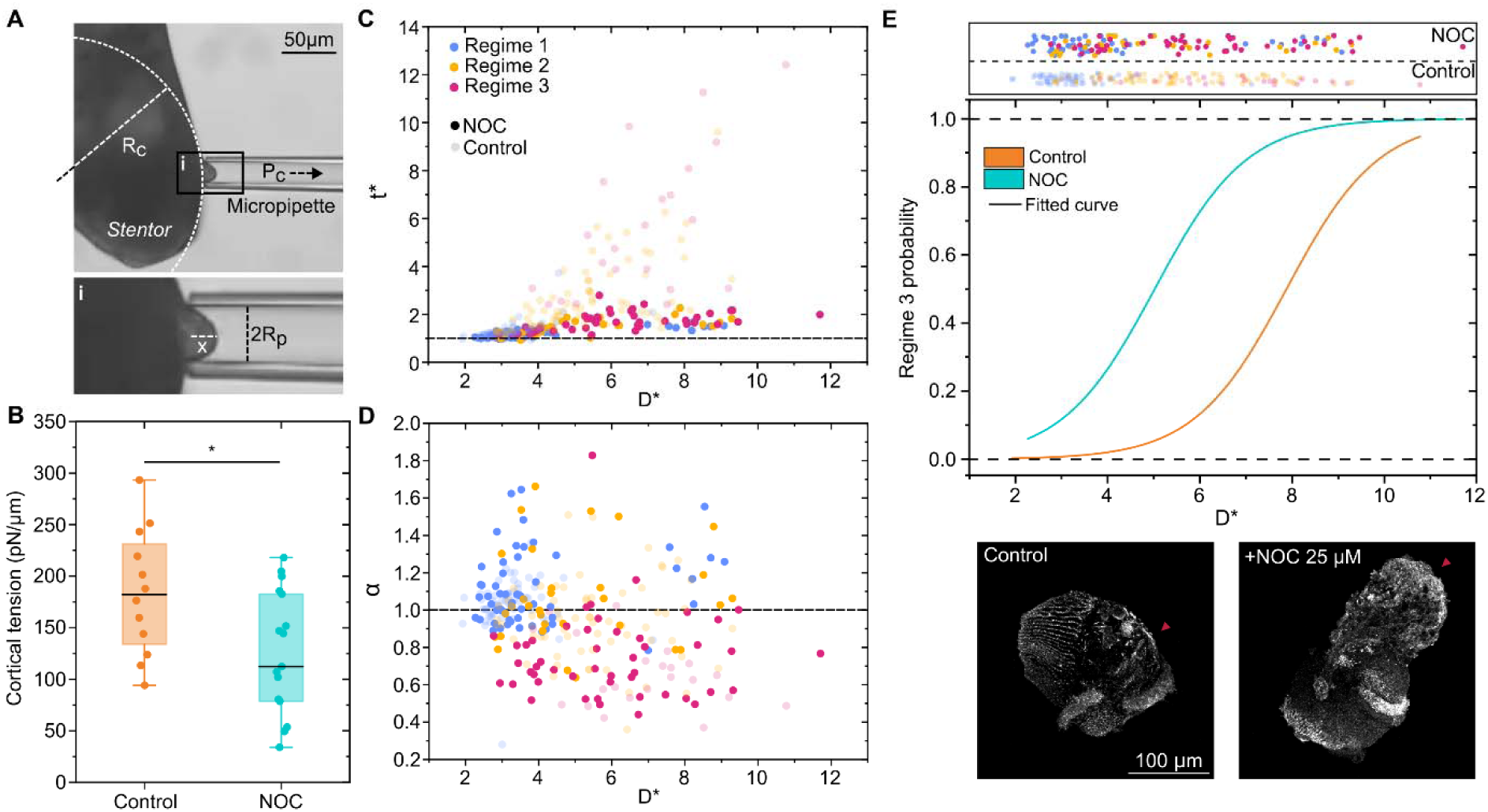
Effects of nocodazole treatment on wounding outcomes. **A.** Micropipette aspiration of a *Stentor* cell using a 20 μm diameter glass micropipette. The inset is a schematic diagram showing the relevant parameters for estimating the cortical tension of the cells. **B.** Plot comparing the cortical tension measured in untreated control cells and the nocodazole-treated cells (p < 0.05). **C.** Plot showing the dimensionless transit time t* and the dimensionless cell size D* for nocodazole-treated cells for the three wounding regimes, overlaid on top of the data (half transparent) for untreated control cells **D.** Plot showing the deformation parameter α and the dimensionless cell size D* for nocodazole-treated cells for the three wounding regimes, overlaid on top of the data (half transparent) for untreated control cells. **E.** Plot showing the probability of regime 3 outcomes for control and nocodazole-treated cells as a function of the dimensionless cell size D*. The curves represent the best fit sigmoid curve corresponding to the data. The black dotted lines represent probability values of 0.0 and 1.0. The panel at the top shows the control and the nocodazole-treated cells plotted as a function of D* for the three wounding regimes. The color scheme for the regimes is the same as the plots in **C** and **D**, where blue, yellow, and red represent wounding regimes 1, 2, and 3 respectively. Solid symbols represent nocodazole-treated cells. Faded symbols represent control cells. The bottom panels show representative immunofluorescence images of wounded cells: untreated control cells (left), and nocodazole-treated cells (right). In both conditions, cells were injected at 4 mL/h through a 70 μm constriction. The red arrow indicates the sites of wounds.

Next, we injected the nocodazole-treated cells into the microfluidic constriction at a fixed flow rate (4 mL/h). We made three observations. First, across regimes 1, 2 and 3, the normalized transit time t* through the constriction was shorter in nocodazole-treated cells than in control cells (Fig. 3C and ESI Fig. S5A). The cortical tension and/or stiffness of the cell is known to directly influence the pressure gradient required for the transit through a microfluidic constriction.^42^ In addition to reducing cortical tension, nocodazole may also disrupt intracellular microtubules and decrease the cell’s bulk stiffness, although we do not have direct measurements of intracellular microtubules due to limitations of our current immunostaining protocol. The reduced cortical tension in nocodazole-treated cells rendered the cells more deformable and allowed the cell to squeeze through the constriction at a lower pressure gradient across the constriction when there was an occlusion at the constriction, thereby shortening the transit times.

Second, the ability of the cells to recover their shape after transiting through the constriction, as measured by the parameter *α*, was only slightly different in nocodazole-treated cells vs. control cells (Fig. 3D and ESI Fig. S5B). The reduced cortical tension in the treated cells impaired their ability to resist deformation (i.e., treated cells became more deformable), resulting in slightly higher α values in regimes 1 and 2. Because most cells already spanned almost the full width of the channel downstream of the constriction, further increase in *α* was not observable in our experiments due to this width confinement. In regime 3, once a catastrophic membrane rupture occurred, α was dominated by cytoplasmic spillage. As a result, cortical tension—regardless of its magnitude—had little effect on α, and there was no observable difference in α between nocodazole-treated cells and control cells.

Third, the probability of regime 3 outcomes (i.e., rupture wounds) was higher in nocodazole-treated cells than in control cells (Fig. 3E). We focused on regime 3 because it measures the critical failure of the membrane and the cytoskeleton. A sigmoidal fit to the binary outcomes of non-rupture (regimes 1 and 2) vs. rupture (regime 3), using the function 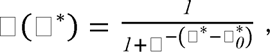 generates 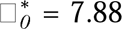 for control cells and 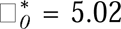 for nocodazole-treated cells, respectively. Here, 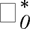 indicates the dimensionless cell size at which the probability *P* of rupture is 50%. This reduction in 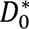 suggests that nocodazole-treated cells ruptured at smaller cell sizes than control cells. The lower cortical tension in nocodazole-treated cells likely reflects a reduction in cortical stiffness, which compromised the cell’s ability to resist large mechanical stresses and increased the likelihood of the failure of the cortex resulting in rupture wounds.

Based on the cortical tension measurements, we estimated the cortex capillary number 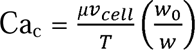 to be in the range 0.78 – 2.87 for nocodazole-treated cells and 0.49 – 1.93 for control cells. Capillary number represents the ratio of viscous forces to cortical tension. The term 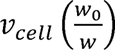 is the estimated average flow velocity at the constriction. v_cell_ is the cell velocity in the straight part of the channel where the width is w_0_, and w is the width of the constriction. μ = 10^-3^ Pa s is the viscosity of the cell medium. T is the cortical tension. The higher capillary number in nocodazole-treated cells indicates that viscous forces, which drive cell deformation, became more dominant relative to cortical tension, which resists deformation. This shift led to greater cell deformation and potentially increased the likelihood of membrane rupture. Together, these results indicate that nocodazole-induced weakening of the KM fiber network reduced the cell transit time and rendered the cells less resistant to membrane rupture.

### Wound resistance increases until a critical hydrodynamic shear rate

Thus far, our experiments were carried out at a constant flow rate. However, we expect the flow rate to influence the probability of membrane rupture, as it governs the hydrodynamic stresses exerted on the cells. Therefore, we varied the flow rate from 2 – 23 mL/h (with the corresponding Ca_c_ from 0.18 to 12). Surprisingly, some large cells with high D* values that were typically wounded (regime 3) at low Ca_c_ remained unwounded (regime 1) at high Ca_c_ (Fig. 4A). Additionally, the decision boundary—defined as the contour corresponding to a regime 3 probability of 0.1—initially exhibited a positive slope in the Ca_c_ vs. D* plot, indicating that cells at higher Ca_c_ had increased resistance to rupture. This upward trend persisted up to a threshold around Ca_c_ ∼ 3. Beyond this point, the decision boundary reversed direction and acquired a negative slope, suggesting that further increases in Ca_c_ were associated with a higher likelihood of rupture.

**Fig. 4.**
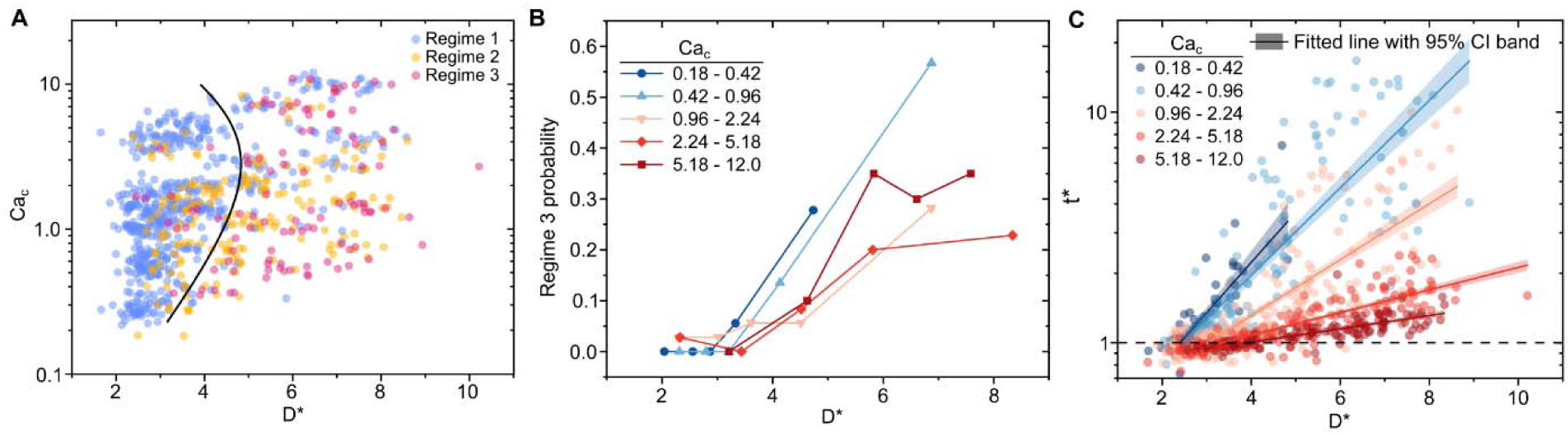
Wound resistance under large hydrodynamic stress. **A.** Regime map showing the rupture (regime 3) and non-rupture (regimes 1 and 2) outcomes as functions of the cortex capillary number Ca_c_ and the dimensionless cell diameter D*. The curve in black denotes the decision boundary drawn at a rupture (regime 3) probability prediction of 0.1 based on logistic regression of the outcomes. **B.** Cumulative probability distribution of regime 3 outcomes (i.e., rupture wounds) as a function of the dimensionless cell size D* for ranges of the cortex capillary numbers Ca_c_. **C.** Plot showing the dimensionless transit time t* as a function of the dimensionless cell size D* for different ranges of the cortex capillary number.

This observation is consistent with the cumulative distribution of the binary outcomes of non-rupture (regimes 1 and 2) vs. rupture (regime 3) at different Ca_c_ (Fig. 4B). Although the binning was coarse (see Materials and Methods), the data shows that increasing Ca_c_ initially shifted the rupture probability curves rightward, suggesting cells with a given D* were less likely to be ruptured at higher Ca_c_ values. Beyond a threshold of Ca_c_ ∼ 2.24 - 5.18, this trend started to be reversed. These results are in contrast to the hydrodynamic stability of droplets and capsules which show monotonically increasing likelihood of rupture on increasing the capillary number.^59,67^

We set out to identify potential physical interpretations that could reconcile this paradoxical observation. We found that the dimensionless transit time t*, plotted for all regimes, decreased and approached t* = 1 as the cortex capillary number increased (Fig. 4C). From the definition of t*, t* = 1 corresponds to the flow of the continuous phase at the constriction. This observation suggests that at higher cortex capillary numbers, the cells exhibited flow behavior resembling that of the continuous phase as they passed through the constriction. We note that some data with t* < 1 arose from insufficient frame rate and cell deformation during the transit through the constriction. Previously, it has been found that the transit time of capsules through constrictions is correlated to their mechanical properties.^45,68,69^ Specifically, the transit time decreases with decreasing viscosity ratio between the capsule and the continuous phase.^54^ In certain cell types, including amoeba,^70,71^ *E-coli* extract^72^ and neutrophils,^68,73^ the cytoplasm is known to exhibit shear thinning behavior. Shear thinning is common in systems where shear forces might cause the breakdown of aggregates (such as protein polymers).^74,75^ In amoeba, the active remodeling of the cytoskeleton^70^ and the sol-gel conversion of the actomyosin network^71^ are responsible for the shear-thinning property of the cell cytoplasm.

Therefore, it is possible that the cytoplasm of *Stentor* cells also exhibits shear-thinning behavior. To attempt to characterize the rheological behavior of *Stentor* cells, we measured the shear rate dependence of the viscosity of a cell suspension. Direct measurement of the cell lysate was not practical in *Stentor* because of the low density of cells in culture (typically less than 100 cells/mL). Additionally, the high cilia-mediated motility prevented in-situ live-cell characterization such as atomic force microscopy. Our rheological characterization of the cell suspension showed that for shear rates in the range of 100 s^-1^ to 5000 s^-1^, the viscosity of the samples decreased as the shear rate increased (ESI Fig. S6). We chose this range of shear rates to match the range of shear rates that the cells experience inside our microfluidic device (∼ 45 s^-1^ to 2800 s^-1^ in the straight part of the channel). We do not believe that the shear-thinning behavior observed was caused by the interaction between the cells^76,77^ because we used a low volume fraction of cells in the suspension (≤ 0.12) ^76^ and we used a thin gap between the parallel plates in the rheometer to ensure that there was one layer of cells in the gap.^77^

Although the shear-thinning rheology of the cells may enhance their ability to resist wounds at elevated cortex capillary numbers, beyond a critical threshold, hydrodynamic stresses exceed their capacity to resist wounds, resulting in the observed reversal in the trend of the decision boundary in Fig. 4A. Finally, to collapse the data from nocodazole treatment with the data from increased hydrodynamic stress in a single regime map would likely require more accurate measurements of the cell viscosity. The phase diagram of rupture vs. no rupture events might be better represented by a parameter combining the capillary number and the viscosity ratio of the cells to the continuous phase. However, this analysis is beyond the scope of the current study due to challenges in obtaining precise cell viscosity measurements.

## Conclusions

In summary, this study has provided an initial framework for characterizing cellular wound resistance in *Stentor.* We investigated the conditions under which *Stentor* became wounded. Larger cells had longer transit times through a microfluidic constriction and were more susceptible to rupture compared with smaller cells. They also exhibited impaired shape recovery after transit due to membrane damage and cytoplasmic loss. Weakening of the *Stentor* KM fiber bundles compromised the cell’s ability to withstand mechanical stresses inside the microfluidic constriction and increasing the likelihood of membrane rupture. We also observed that cells were more resistant to higher hydrodynamic stress up to a threshold, likely due to the shear thinning of the cell cytoplasm. Together, our findings suggest that *Stentor* leverages both its internal cytoskeletal architecture and the rheological properties of its cytoplasm to resist mechanical stress, revealing a robust cellular strategy for wound resistance in the absence of rigid external structures.

While prior work in soft matter has explored deformation and rupture in synthetic capsules and vesicles, *Stentor* represents a living, capsule-like system with active biophysical responses. Our findings highlight both similarities and key differences between engineered soft materials and biological systems, particularly the contributions of the microtubule bundles and shear-thinning cytoplasm in resisting wounding. This study expands the concept of mechanical robustness from passive soft materials to actively maintained living systems, provides a framework for exploring wound resistance in single-celled organisms lacking an external shell, and introduces a microfluidic platform for probing cellular mechanics under stress.

For future work, a more detailed examination of the mechanical properties of *Stentor* cells will be necessary to build upon the findings in this study. Because *Stentor* is highly motile, the use of conventional mechanical characterization methods such as atomic force microscopy is not feasible. *Stentor* also has a trumpet shape, making it challenging to directly apply certain microfluidic models developed for capsules or mammalian cells to extract its mechanical properties. Further investigation is also needed to clarify the mechanical role of the KM fibers. The treatment of *Stentor* cells with the microtubule stabilizing drug taxol did not show significant difference in wound resistance when compared with untreated cells (Fig.S7, ESI). We attribute this observation to the fact that KM fibers are stable structures composed of acetylated microtubule bundles, and further stabilization by taxol is unlikely to significantly affect wound resistance. Future investigation should also explore the role of the myoneme network. Finally, the microfluidic system as a generalized tool has the potential to be useful for comparing wound resistance across diverse single-celled organisms to further elucidate the role of cellular structures and rheological properties in conferring wound resistance.

## Data availability

Data pertaining to this study will be made available upon reasonable request.

## Acknowledgment

The work was supported by the National Science Foundation (NSF Award 2317442), and in part by the Center for Cellular Construction, which is a Science and Technology Center funded by the National Science Foundation (NSF Award: DBI-1548297). WFM acknowledges support from NIH grant R35 GM130327. Device fabrication was performed in the Stanford Nano Shared Facilities (SNSF) and the Stanford Nanofabrication Facilities (SNF). Imaging was carried out at the Cell Sciences Imaging Center (CSIF) at the Stanford School of Engineering Shriram Center. We would like to acknowledge members of the Tang and Marshall labs for helpful discussions. Special thanks to the Moumita Das lab at Rochester Institute of Technology and Manu Prakash at Stanford University for their thoughts on the project.

