## Supplementary Information for "Characterization of cellular wound resistance in the giant ciliate *Stentor coeruleus*"

**Electronic Supplementary Information**

**Fig. S1.** **Microfluidic channel designs. A.** The design for injecting cells at low flow rate (≤ 10 mL/h). The inset shows the microfluidic constriction. **B.** The design for injecting cells at high flow rate (> 10 mL/h). It consists of a primary inlet for cell injection and a secondary inlet for sheath flow.

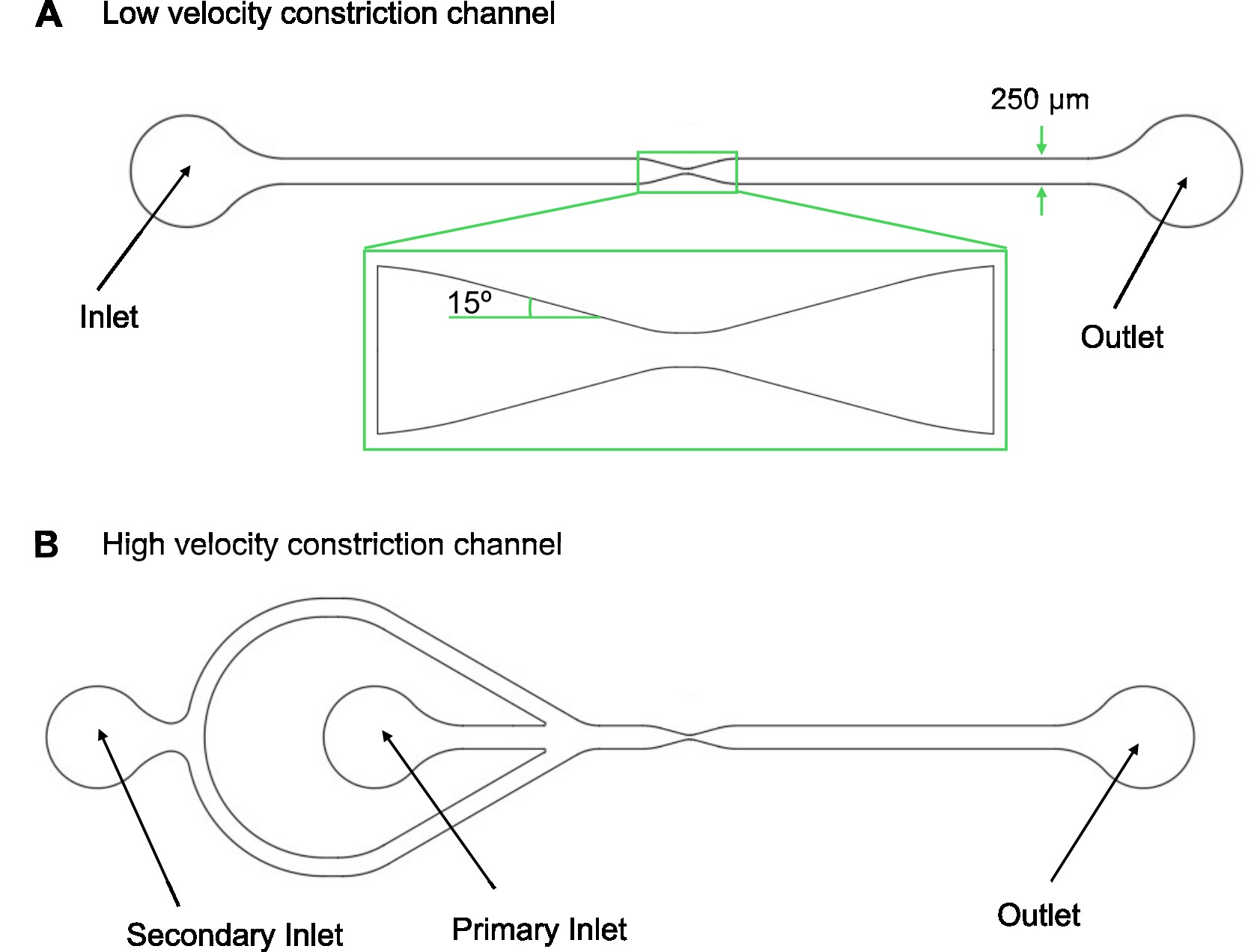

**Fig. S2.** Schematic diagram for estimating the characteristic timescale for cell transit. The characteristic timescale is estimated as the time taken by a fluid of the same volume as the cell to pass through the constriction. First, the time taken for the cell to enter the region of interest (ROI) is given by $\tau_{1}=\frac{l_{cell}}{v_{cell}}$. Next, the time taken by the cell to traverse through the ROI is given by $\tau_{2}=\frac{Q_{ROI}}{\dot{Q}}$ where Q_ROI_ is the volume enclosed by the ROI and $\dot{Q}$ is the flow rate. On simplification, we get, $\tau_{2}=\frac{A_{ROI}}{v_{cell}w_{0}}$. Hence, the characteristic transit time is given by:

$$\tau=\tau_{1}+\tau_{2}=\frac{l_{cell}}{v_{cell}}+\frac{A_{ROI}}{v_{cell}w_{0}}$$

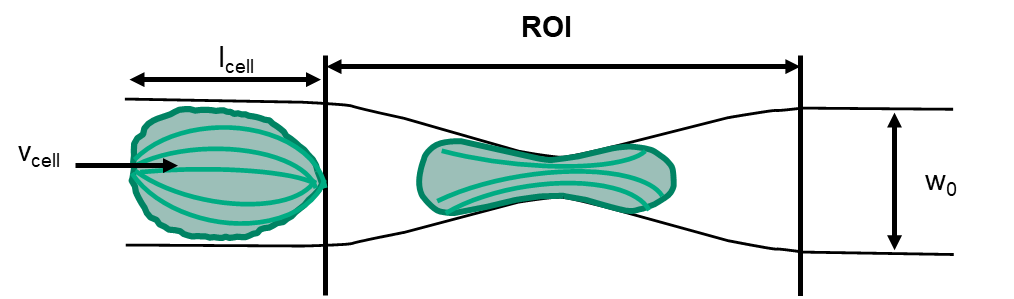

**Fig. S3. Examples of regime 2 outcomes.** **A.** Membrane shearing in the upstream part of the cell. **B.** A bleb forms usually in the downstream part of the cell.

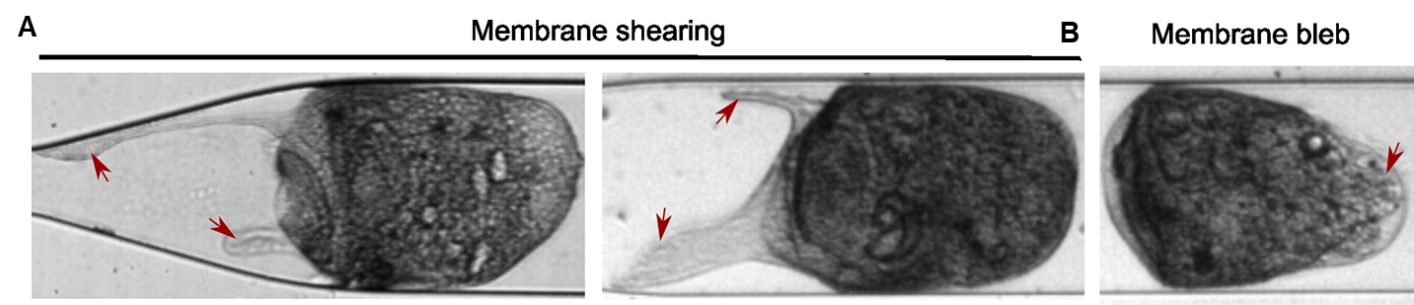

**Fig. S4.** Immunofluorescence images of the KM fiber network in control cells and nocodazole-treated cells. The red arrows show the unusual puncta in the nocodazole-treated cells. Brightness and contrast have been enhanced for ease of visualization.

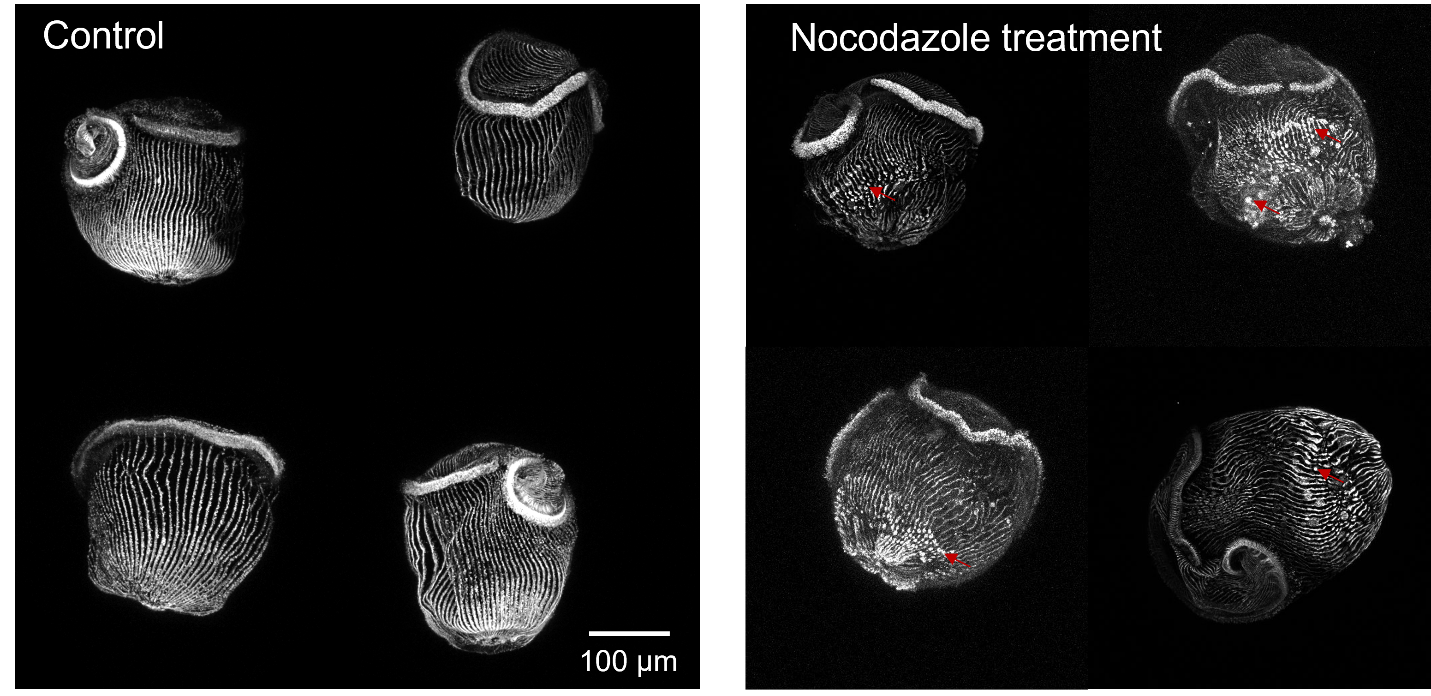

**Fig. S5. A.** Plot comparing the dimensionless transit time t* for the three wounding regimes between untreated control cells and nocodazole-treated cells. **B.** Plot comparing the deformation parameter α for the three wounding regimes between untreated control cells and nocodazole-treated cells.

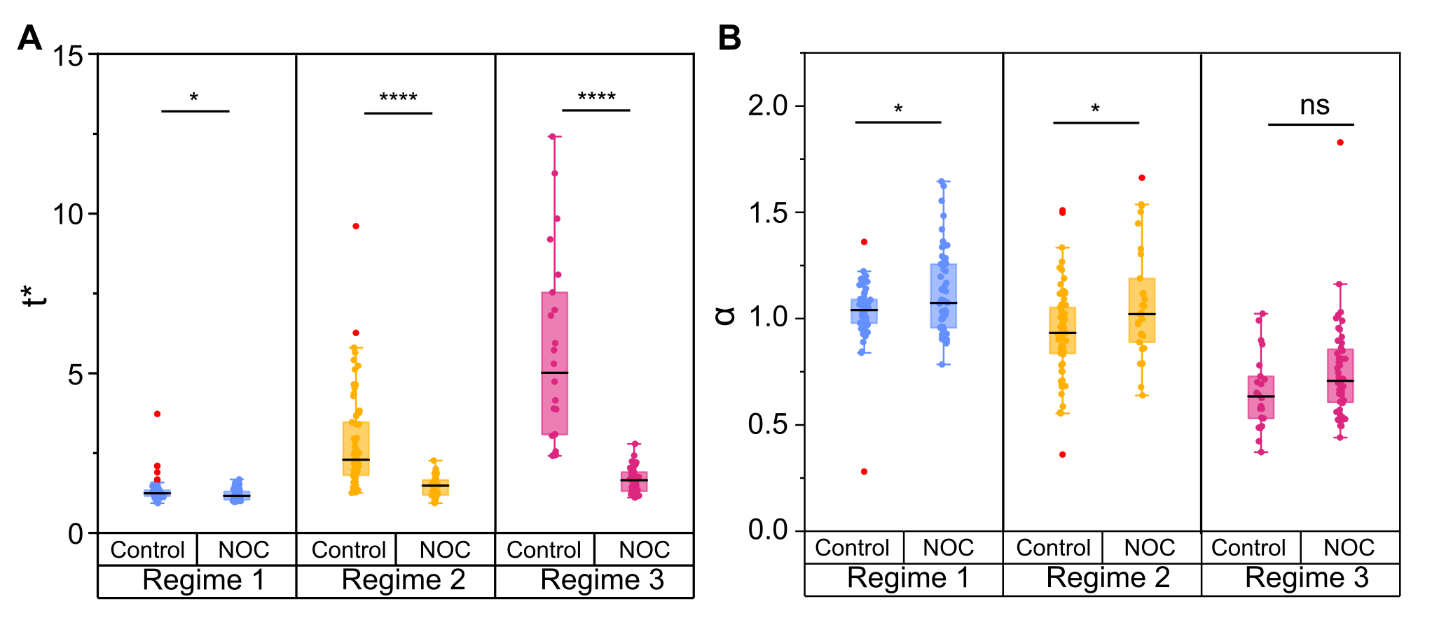

**Fig. S6.** Plot showing the shear rate dependent viscosity of cell samples of varying cell volume fractions. The inset shows a schematic diagram of the parallel plate configuration of the rheometer used in the characterization. The error bar represents one standard deviation from the mean of three sets of measurements, each consisting of N~300 cells (for cell volume fraction $\phi$ = 6%) and N~600 cells ($\phi$ = 12%), respectively. Each set of measurements used a new population of cells.

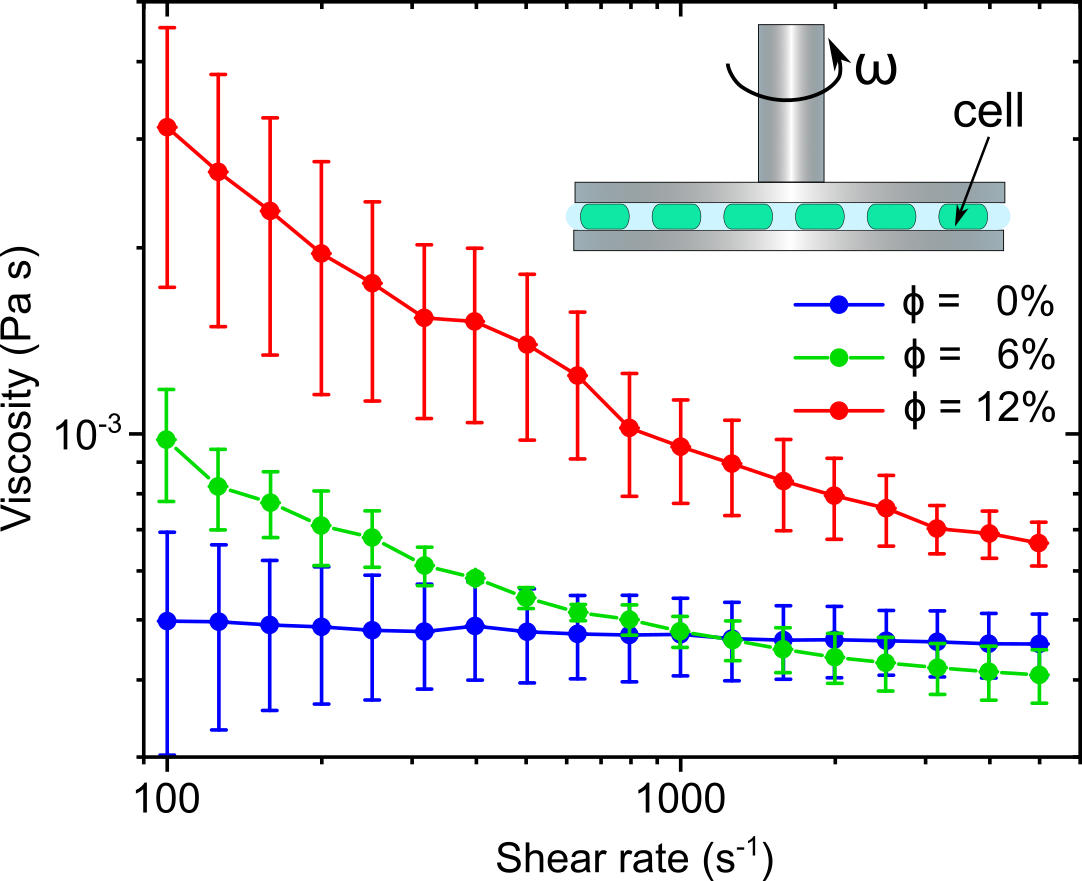

**Fig. S7. Effect of taxol on experimental outcomes. A.** Plot comparing the dimensionless transit time t* for the three wounding regimes between untreated control cells, nocodazole-treated cells and taxol-treated cells. **B.** Plot comparing the deformation parameter α for the three wounding regimes between untreated control cells, nocodazole-treated cells, and taxol-treated cells. **C.** Plot showing the probability of regime 3 outcomes for control, nocodazole-treated and taxol-treated cells as a function of the dimensionless cell size D*. The curves represent the best fit sigmoid curve corresponding to the data. The black dotted lines represent probability values of 0.0 and 1.0. The panel at the top shows the distribution of the regimes 1 (blue), 2 (yellow) and 3 (red) data points for the three conditions.

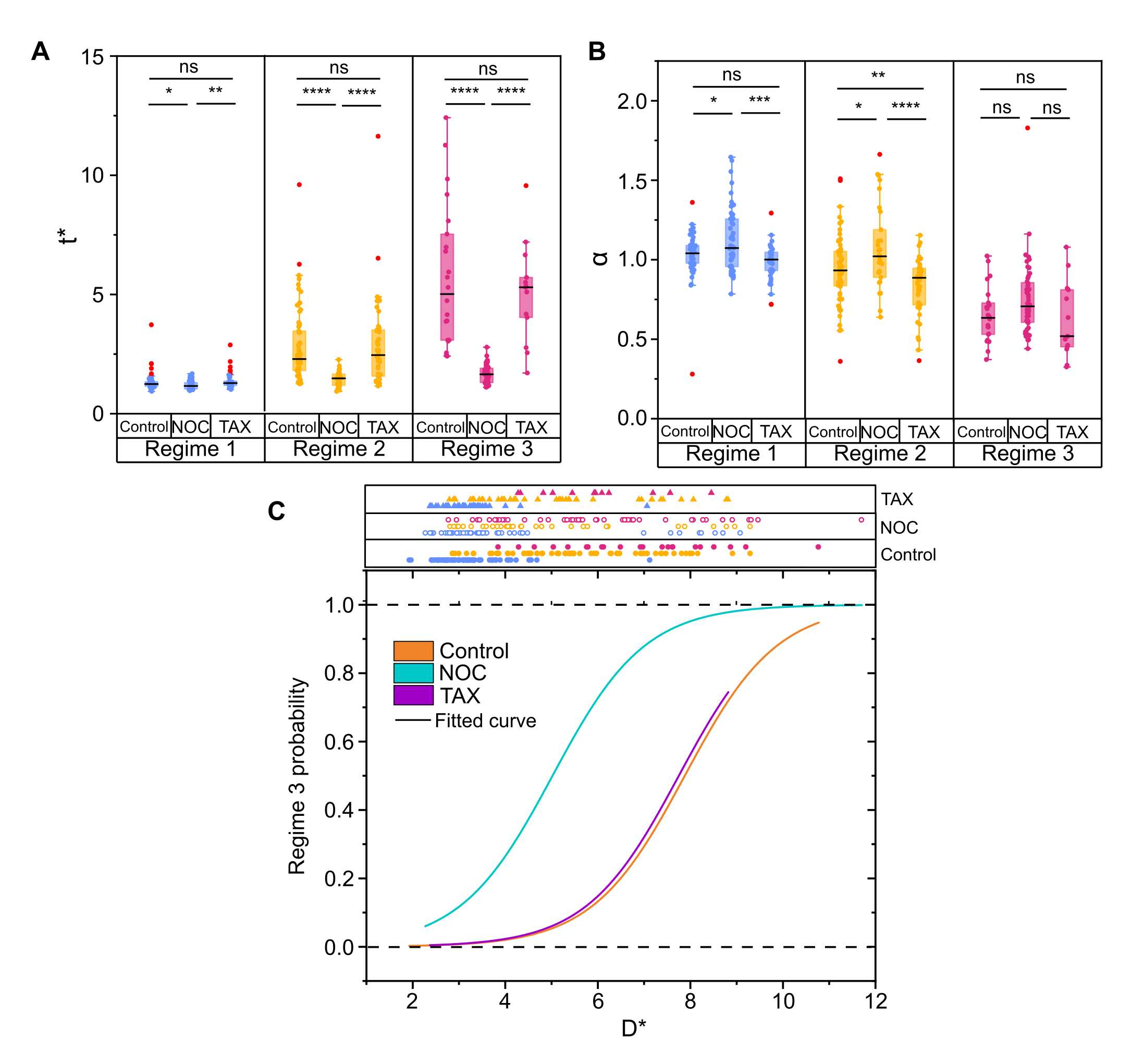
